# Distinct trans-placental effects of maternal immune activation by TLR3 and TLR7 agonists: implications for schizophrenia risk

**DOI:** 10.1101/2021.09.20.460754

**Authors:** Jaedeok Kwon, Maria Suessmilch, Alison McColl, Jonathan Cavanagh, Brian J. Morris

## Abstract

Exposure to infection *in utero* predisposes towards psychiatric diseases such as autism, depression and schizophrenia in later life. The mechanisms involved are typically studied by administering mimetics of double-stranded (ds) RNA viral or bacterial infection to pregnant rats or mice. The effect of single-stranded (ss) virus mimetics has been largely ignored, despite evidence linking prenatal ss virus exposure specifically with psychiatric disease. Understanding the effects of gestational ss virus exposure has become even more important with the current SARS-CoV-2 pandemic. In this study, in pregnant mice, we compare directly the effects, on the maternal blood, placenta and the embryonic brain, of maternal administration of ds-virus mimetic poly I:C (to activate toll-like receptor 3, TLR3) and ss-virus mimetic resiquimod (to activate TLR7/8). We find that, 4h after the administration, both poly I:C and resiquimod elevated the levels of IL-6, TNFα, and chemokines including CCL2 and CCL5, in maternal plasma. Both agents also increased placental mRNA levels of IL-6 and IL-10, but only resiquimod increased placental TNFα mRNA. In foetal brain, poly I:C produced no detectable immune-response-related increases, whereas pronounced increases in cytokine (e.g. *Il-6, Tnf*α) and chemokine (e.g. Ccl2, *Ccl5*) expression were observed with maternal resiquimod administration. The data show substantial differences between the effect of maternal exposure to a TLR7/8 activator as compared to a TLR3 activator. There are significant implications for future modelling of diseases where maternal ss virus exposure contributes to environmental disease risk in offspring.

## Background

Maternal bacterial and viral infections during pregnancy increase the risk of psychiatric disease in the offspring exposed to the infection *in utero*. This is the case for autism spectrum disorders, major depressive disorder (Brown and Meyer, 2018; al-Haddad et al., 2019) and schizophrenia (Brown et al., 2001; Brown, 2006; Brown, 2012). The mechanisms involved remain unclear, but are likely to involve the exposure of the developing foetal brain to either the infectious agent itself, or to cytokines and chemokines released from maternal or placental tissues as part of the innate immune response. Direct exposure of the developing foetal CNS to the infectious agent is possible, as the agents most strongly linked by epidemiological evidence to schizophrenia risk (rubella virus, influenza virus and the parasite *Toxoplasma Gondii*) (Brown et al., 2001; Brown, 2006; Mortensen et al., 2007; Brown, 2012), are all able to penetrate the placenta to invade the foetal environment(Rasmussen et al., 2012; Leung et al., 2019; Ostrander and Bale, 2019). Whatever the mechanism, the infectious agent or immune response molecules impact upon differentiating or migrating neurons e.g inter-neurons, and can affect microglia in the developing CNS with consequences for processes such as synaptic pruning (Wang et al., 2019). Exactly how these processes operate to raise risk of psychiatric disease remains uncertain.

Maternal immune activation models typically involve pregnant mice being injected with the bacterial mimetic lipopolysaccharide (LPS), or the double-stranded (ds) virus mimetic polyinosinic: polycytidylic acid (poly I:C). The consequences for the neurochemical and behavioural phenotype of the offspring can then be evaluated. A variety of disease-relevant changes in gene expression and social, affective and cognitive behaviours have been detected in offspring at various ages (Meyer et al., 2009; Meyer, 2014; Estes and McAllister, 2016), although there have also been concerns about the reproducibility of some of the data with poly I:C, due to a variety of technical factors (e.g. batch variation, endotoxin contamination) (Kowash et al., 2019; Mueller et al., 2019).

LPS reproduces the effects of bacterial infection by stimulating the toll-like receptor (TLR) TLR4, while poly I:C reproduces the effects of ds-virus infection by stimulating TLR3. Ss-viruses stimulate TLR7 or TLR8 to induce an immune response. However, despite the fact that some of the prenatal infectious agents most strongly linked to schizophrenia risk (rubella virus and influenza virus) are ss-viruses, models of maternal immune infection with ss-virus mimetics such as resiquimod or imiquimod, which stimulate TLR7/8 (Hemmi et al., 2002; Isobe et al., 2006) are rarer. Similarly, *Toxoplasma Gondii* also induces an innate immune response via TLR7 (Andrade et al., 2013; Yarovinsky, 2014), while ds-viruses such as Herpes simplex II are not robustly linked to schizophrenia risk (Brown et al., 2006). Furthermore, some recent evidence in mice suggests that the behavioural consequences for offspring exposed to TLR7 stimulation *in utero* may be different to the effects of maternal TLR3 or TLR4 stimulant exposure (Missig et al., 2020).

Our research question in this study was: what are the differences in the cytokine/chemokine responses to the commonly-used ds viral mimic versus ss viral mimic. We have directly compared the most commonly used strategy for studying the effects of MIA (poly I:C administration in mice) with a novel strategy using resiquimod to mimic ss-virus infection.

## Methods

### Experimental design

Poly I:C is frequently used in mice to model MIA. Concerns have been raised recently about variability in its effects, due partly to possible endotoxin contamination and batch variation (Kowash et al., 2019; Mueller et al., 2019). The dose used is most commonly 20mg/kg, and the gestational time point is generally E9, E12.5 or E17, with E12.5 the most frequent choice, as it is argued to correspond in various ways to mid-gestation time in humans (Meyer et al., 2009; Meyer, 2014; Estes and McAllister, 2016; Brown and Meyer, 2018). We used poly I:C of the specification that reportedly gives the most robust results (Kowash et al., 2019; Mueller et al., 2019), and selected resiquimod as the ss-virus mimetic comparator. While imiquimod has recently been used to study the consequences for CNS function of ss-virus infection (Thomson et al., 2014; McColl et al., 2016), and to assess the behavioural effects in offspring of MIA (Missig et al., 2020), apart from acting as a ss-virus mimetic, imiquimod also has direct actions on adenosine receptors that complicate interpretation of its effects (Schön et al., 2006; Wolff et al., 2013). In addition, imiquimod is selective for TLR7, whereas resiquimod activates both TLR7 and TLR8 (Patinote et al., 2020). Considering that the epidemiological evidence for psychiatric disease risk implicates maternal ss-virus infection, without discriminating between TLR7 and TLR8 mediation, resiquimod seems to be the better choice to maximise construct validity in future models of environmental contribution to psychiatric disease risk.

Resiquimod has been used systemically in mice on occasion, producing an immune response at doses between 1 and 10 mg/kg (Su et al., 2005; Baenziger et al., 2009; Adzavon et al., 2017; Michaelis et al., 2019). McAllister et al (McAllister et al., 2013) describe equivalent (and substantial) serum TNFα responses to high doses of 30mg/kg poly I:C and 10 mg/kg resiquimod in mice. We therefore selected 2 mg/kg as a moderate dose of resiquimod to compare with the standard dose of 20mg/kg poly I:C.

#### Maternal immune activation and sample collections

Wild type mice (C57/BL6, female, aged wk4) were purchased from Envigo. Mice were time mated separated following day, and if the pregnancy was confirmed the day was given as E0. Female mice were weighed and monitored for 12 days. All mice were aged 7-9 weeks and weighed 30+/-1 g at the point of experiment (12.5 days pregnant). In order to induce maternal immune activation (MIA), three treatments, vehicle (PBS, 2 ml/kg, Gibco 14190144), poly I:C (20 mg/kg of a 10 mg/ml solution in PBS, LMW, Invivogen tlrl-picw), resiquimod (2 mg/kg of a 1 mg/ ml solution in PBS, Invivogen tlrl-r848), were administered subcutaneously on E12.5 between 9:00-11:00 am and their conditions were monitored to ensure that there was no sign of any severe sickness symptoms or abnormalities.

4 hours following injections, the pregnant dam was culled with the CO_2_ euthanasia. The maternal blood was collected via right atrium into an EDTA-coated syringe. The blood was injected into an EDTA-coated 1.5 micro centrifuge tube containing 100 ul EDTA and shaken. Experimental samples were collected. Following centrifugation at 10,000g at 4°C for 10 minutes, the supernatant (plasma) was frozen at -80°C for further use. Once the blood collection was done, cut right atrium and PBS perfusion was carried. After the PBS perfusion, maternal placentae and embryos were dissected and collected. All tissues were pre-processed and stored until further uses. More details of sample preparation procedures are described in separate sections. All experiments were reviewed by the University of Glasgow ethical committee and were performed under the authority of UK Home Office License.

#### RT-qPCR

Collected tissues after MIA experiment were originally stored in RNAlater at -80 °C. Before extracting total RNA from the tissues, they had to be completely thawed in RT and RNAlater had to be removed. Under RNase-fee condition, the collected tissues were homogenised using the TissueLyser LT (Qiagen) in 600 μl of prepared RLT lysis buffer (Qiagen). Total RNA was extracted from the stabilised tissues using an RNeasy mini kit (Qiagen, 79254) with additional DNase I (Qiagen, 79254) as per manufacturer’s instruction. The RNA concentration was determined by a Nanodrop DeNovix DS-11+Spectrophotometer. RNA was reverse-transcribed to cDNA using High-capacity RNA-to-cDNA™ kit (Applied Biosystems, 4387406) according to the manufacturer’s instruction. cDNA concentration was normalised by using equal quantity of RNA.

Gene expression in the tissues was quantified Fast SybrGreen™ master mix (Applied Biosystems, 4385612) for each target using the QuantaStudio7 (Thermo Fisher Scientist). Samples were run in triplicate on 384 well qPCR plates (Applied Biosystems, 4309849) and the levels of the target genes were normalised to a geometric mean of two housekeeping genes (*Gapdh, Tbp*) and relative differences in target gene expression were determined using absolute quantification method. Primer sequences are provided in Supplementary Table 2.

#### Multiplex/Luminex

The concentration of 9 cytokines and chemokines with well-characterised roles in innate immune responses were measured in maternal plasma by a Multiplex/Luminex (Merck, MCYTMAG-70K-PX32) according to the manufacturer’s instruction. Measurements were taken (LUMINEX 200^®^), operated via Bio-Rad’s Bio-Plex software version 6.1™. The beads were read determining the mean fluorescence intensity (MFI) (Breen et al., 2015; Breen et al., 2016; Richter et al., 2017) and the data were accepted if the duplicated samples vary (CoV) were less than 15%.

#### Statistical Analysis

All statistical analyses were carried out using Minitab 19 Statistical Software and all data were reported as group mean ± S.E.M. Luminex data were Box-Cox transformed prior to one-way analysis of variance (ANOVA) with post hoc Fisher LSD tests to correct for multiple comparisons. Non-normally distributed RT-qPCR data were log transformed, followed by two-way ANOVA with post Tukey tests. A p-value of <0.05 was considered significant.

## Results

### Protein level changes in maternal plasma

Following poly I:C and resiquimod there were increases in the levels of immune molecules in maternal plasma, compared to vehicle administration. Of the 9 cytokine/ chemokines tested, IL-6, TNF, IL-10, CCL2, CCL5, CXCL10, and LIF (leukaemia inhibitory factor) were all elevated following both poly I:C and resiquimod administration: (Figure 1). Maternal plasma from mice injected with resiquimod, but not poly I:C, also showed significantly upregulated CCL11 and CXCL1, (Figure 1). In general, the effects of poly I:C showed a greater variability in the magnitude of cytokine/chemokine induction, compared to resiquimod.

**Figure 1.**
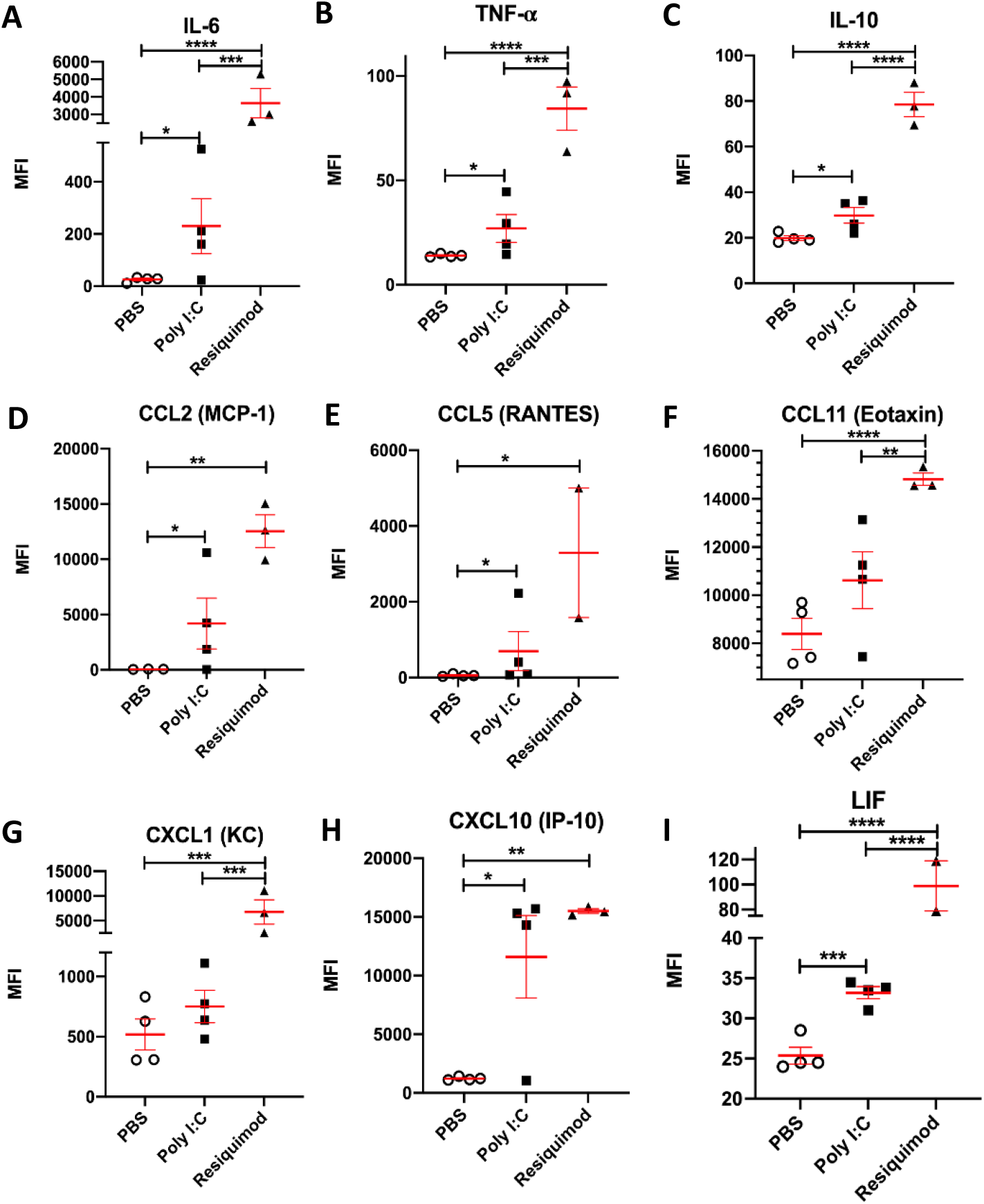
MIA by both poly I:C and resiquimod induces an immune response in maternal plasma. All measured immune molecules were upregulated in the plasma of mothers following resiquimod compared to control (PBS). Poly I:C caused significant upregulation most of measured immune molecules except CCL11 (Eotaxin) and CXCL1 (KC) compared to PBS. The individual dots are shown along with mean ± SEM. The data were box-cox transformed and analysed by one way-ANOVA, Fisher LSD post-hoc test (n=4 independent samples in each condition; *p≤0.05, **p≤0.005, ***p≤0.001, ****p≤0.0001). Details of ANOVA F values and p values are provided in Supplementary Table 1

### Transcription level changes in placentae

The data from maternal plasma clearly confirm that the experimental conditions caused an immune response in the dams. We next assessed the response of placental tissue to the immune stimuli, measuring mRNA levels so that changes detected can be unequivocally ascribed to the placenta, as opposed to maternal plasma.

Resiquimod administration elevated expression of most of inflammatory cytokine and chemokine mRNAs (*Tnf-α, Ccl5, Ccl11* and *Cxcl1*) without affecting *Ccl2* and *Cxcl12* mRNA levels (Figure 2). Poly I:C induced *Il-6, Il-10*, and *Cxcl10* mRNA levels, but changes were not observed in any other genes (Figure 2A, C, and H). While CCL2 in maternal plasma was significantly induced by both poly I:C and resiquimod injection (Figure 1D), placental *Ccl2* mRNA was not changed by either MIA condition compared to vehicle (Figure 2D). Overall, the results suggest that while the response to poly I:C and resiquimod in maternal plasma is rather similar, the response of placental tissue to poly I:C is much more limited compared to the response to resiquimod.

**Figure 2.**
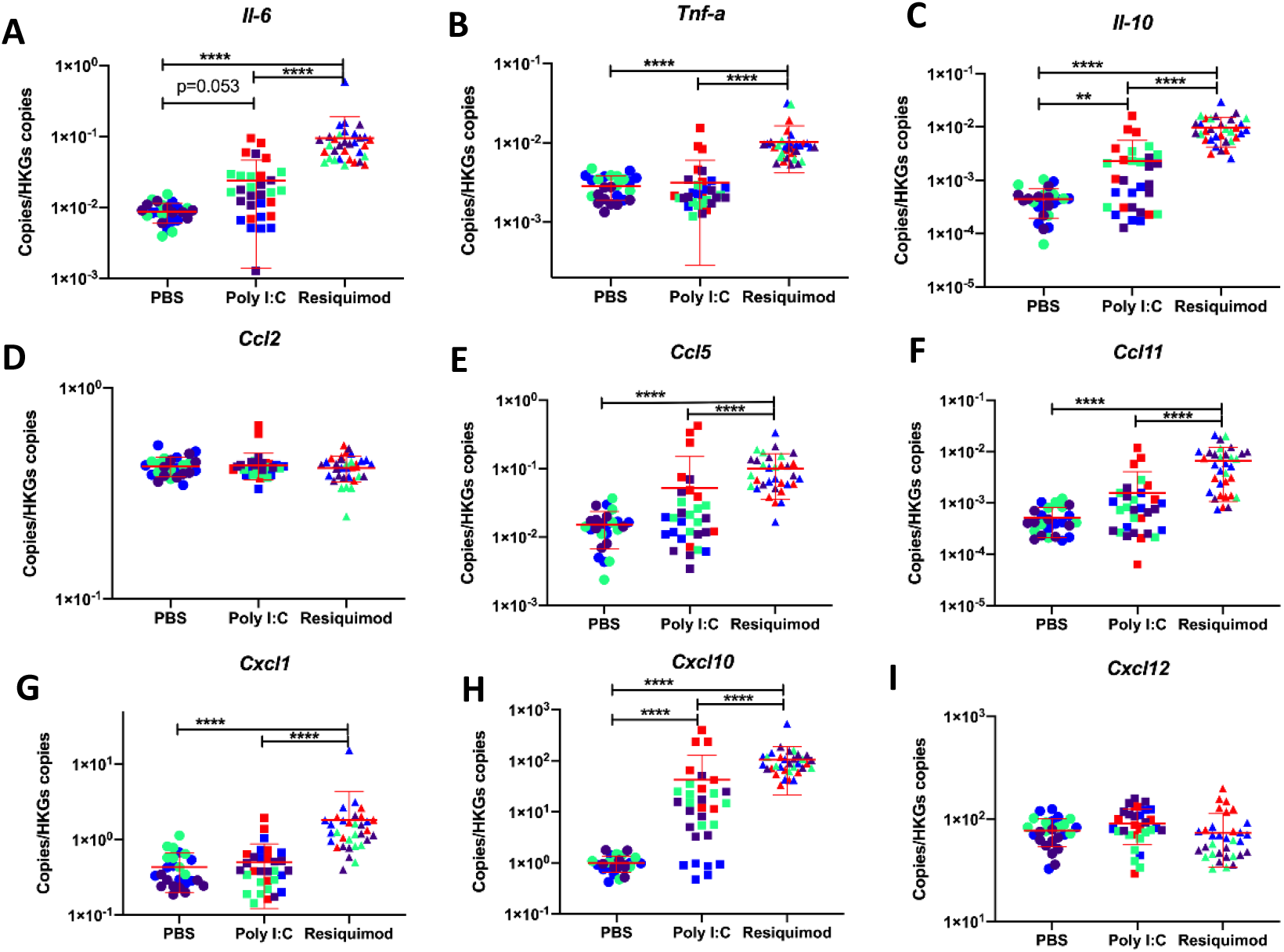
Effect of MIA by poly I:C or resiquimod on immune response in placentae. Placental tissues were collected after 4 hour MIA (PBS, poly I:C (20mg/kg, LMW), resiquimod (2mg/kg)). **(A, C, H)** *Il-6, Il-10*, and *Cxcl10* mRNAs were significantly upregulated by poly I:C and resiquimod compared to PBS. **(B, E, F, G)** *Tnf-a, Ccl5, Ccl11*, and *Cxcl1* were induced by resiquimod, but not by poly I:C compared to PBS. **(D, I)** *CC2* and *Cxcl12* were not changed by MIA. Absolute quantification was performed via RT-qPCR and the data were normalised to *Gapdh* and *Tbp*. The individual dots are shown along with mean ± SEM. Colour indicates dams within a single treatment (same colour means “same dam”). The data were log transformed and analysed by twoway-ANOVA, Tukey post-hoc test (n=26-33 independent samples; *p≤0.05, **p≤0.005, ***p≤0.001, ****p≤0.0001). Details of ANOVA F values and p values are provided in Supplementary Table 1

### Transcription level changes in foetal brains

The same cytokines and chemokines mRNAs were measured in foetal brain tissue 4h after MIA (Figure 3). Foetal brains from MIA caused by resiquimod showed significantly increased levels of most of cytokine and chemokine mRNAs compared to vehicle administration. However, the foetal brain tissue from poly I:C MIA did not show any induction of immune molecule genes.

**Figure 3.**
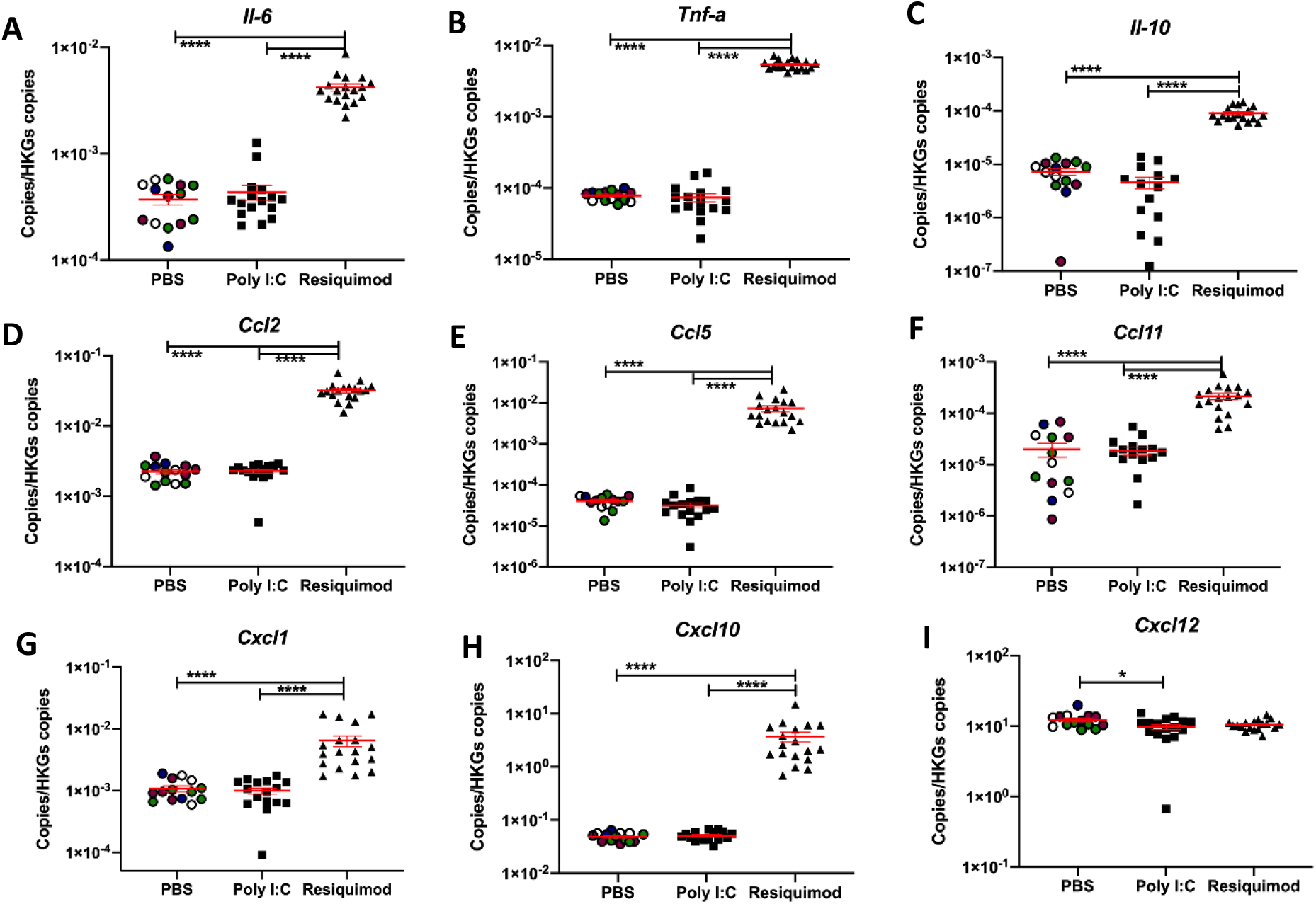
Effect of MIA by poly I:C or resiquimod on immune response in foetal brain. Foetal brain tissues were collected after 4 hour MIA (PBS, poly I:C (20mg/kg, LMW), resiquimod (2mg/kg)). **(A-H)** Cytokine and chemokine mRNAs were significantly induced by resiquimod but not poly I:C compared to control (PBS). **(I)** *Cxcl12* mRNA showed slight changes by MIA. Absolute quantification was performed via RT-qPCR and the data were normalised to *Gapdh* and *Tbp*. The individual dots are shown along with mean ± SEM. The data were log transformed and analysed by twoway-ANOVA, Tukey post-hoc test (n=14-18 independent samples; *p≤0.05, **p≤0.005, ***p≤0.001, ****p≤0.0001). Details of ANOVA F values and p values are provided in Supplementary Table 1

### Immune cell markers

There is considerable evidence for microglial activation in foetal brain following MIA with poly I:C (Smolders et al., 2015; Prins et al., 2018; Murray et al., 2019; Ozaki et al., 2020). We tested possible altered expression of microglial markers in foetal brain tissue after MIA.

The results showed that the expression of *Aif1* (Iba-1) was significantly increased by resiquimod, but not by poly I:C (Figure 4C). In contrast, *Tmem119* and *Cx3cr1* mRNA levels were down regulated by resiquimod (Figure 4A, B). Noticeably, poly I:C caused a significant reduction in the levels of *Tmem119* mRNA, but not *Cx3cr1* mRNA. *Ccr2* mRNA was decreased by both stimuli for MIA, although resiquimod’s impact was greater than poly I:C (Figure 4D). *Ly6c2* mRNA levels were not affected by MIA (Figure 4E).

**Figure 4.**
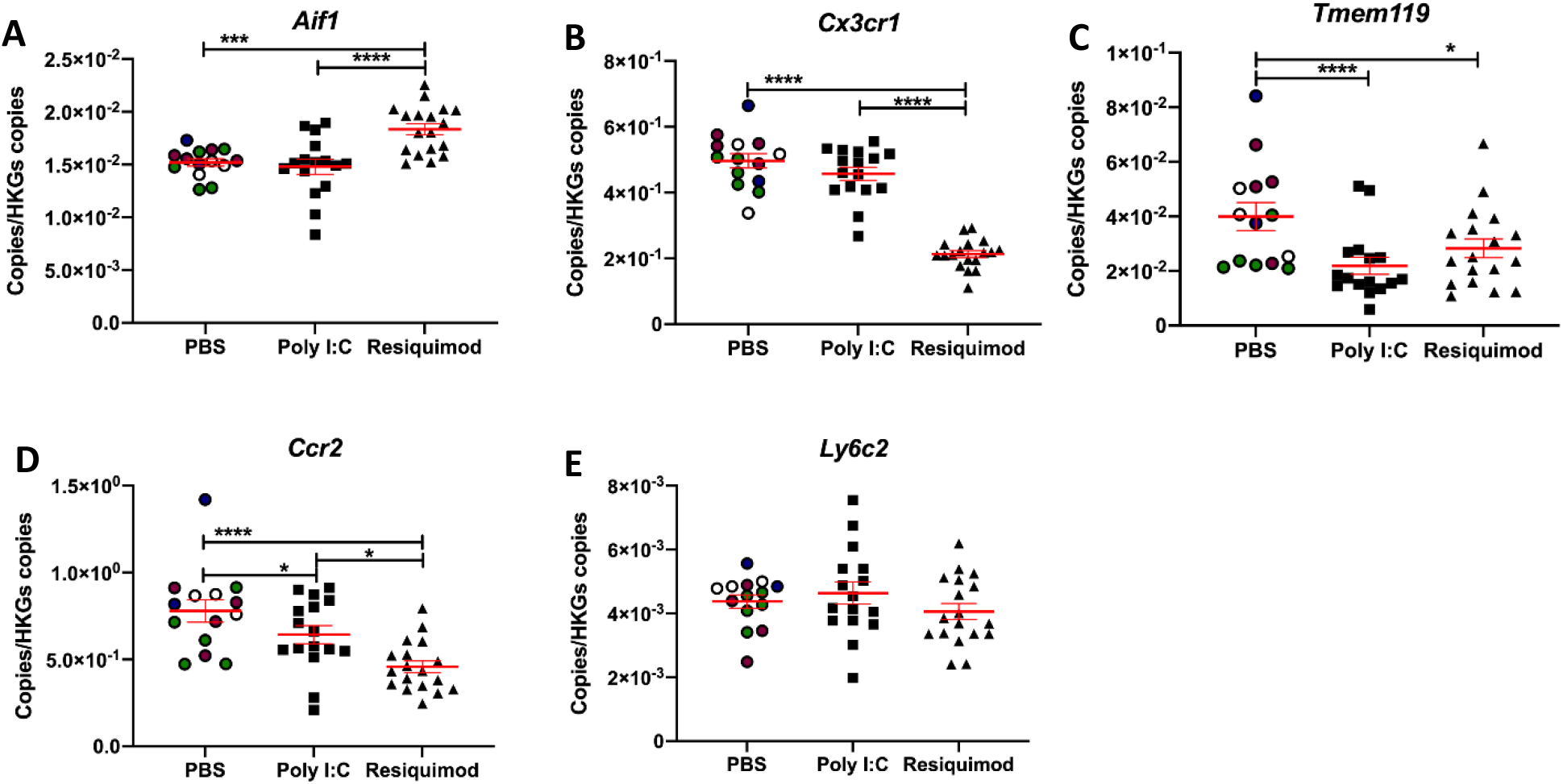
Effect of MIA by poly I:C or resiquimod on microglial markers in foetal brain. Foetal brain tissues were collected after 4 hour MIA (PBS, poly I:C (20mg/kg, LMW), resiquimod (2mg/kg)). **(A**,**B)** *Aif1* was increased by resiquimod, however, *Cx3cr1* was downregulated by resiquimod compared to PBS. Both genes were not changed by poly I:C. **(C)** *Tmem119* was significantly downregulated by MIA compared to PBS. **(D, E)** *Ccr2* were significantly downregulated by MIA compared to PBS, on the other hand, *Ly6c2* was not changed by MIA. Absolute quantification was performed via RT-qPCR and the data were normalised to *Gapdh* and *Tbp*. The individual dots are shown along with mean ± SEM. The data were log transformed and analysed by twoway-ANOVA, Tukey post-hoc test (n=14-18 independent samples; *p≤0.05, **p≤0.005, ***p≤0.001, ****p≤0.0001). Details of ANOVA F values and p values are provided in Supplementary Table 1.

In addition, due to its frequent link to environmental risk for psychiatric disease, we also monitored the expression of *Bdnf*. There were no changes in *Bdnf* expression in foetal brain, but resiquimod exposure decreased *Bdnf* mRNA levels in placenta (Supplementary figure1).

#### Sex effects

Foetal development is affected by sex (Thion et al., 2018; Gildawie et al., 2020). In order to identify any sex-dependent influence on the foetal immune response, the embryos’ sex was determined by qPCR, using the sex-specific *Xist* gene (Cheung et al., 2017). High *Xist* expression indicated female and low expression male (Supplementary Figure2), and suggested that the samples were well-balanced for sex within a treatment group. No statistically significant interaction of sex with the effect of MIA was observed for any of the markers.

## Discussion

This study provides a direct comparison of the pattern of immune response to poly I:C and resiquimod in maternal, placental and foetal brain compartments. It is clear that their patterns of immune activation are different, and the differences are important for future studies of MIA effects on offspring.

### Effects in maternal plasma

Maternal plasma showed evidence of a clear immune response to both poly I:C and resiquimod, with induction of cytokines, including Tnfα and Il-6, and chemokines, including Ccl2 and Ccl5. Maternally-derived Il-6 acting on the placenta, but conceivably originating either in maternal plasma monocytes or in the placenta itself, has been viewed as a key component of MIA effects on the foetus (Hsiao and Patterson, 2011; Wu et al., 2017). Hence the induction of maternal Il-6 observed here is potentially important for some of the downstream events we detect in the foetal brain. However, our data suggested that maternal plasma IL-6 levels were not correlated with mRNA level changes in placenta and embryo brain samples (data not shown). This suggests that the gene induction in placenta and embryo brain samples is less likely to be caused by maternal plasma IL-6.

At the doses used, however, resiquimod tended to produce larger effects than poly I:C. The variability of the maternal immune response to poly I:C in mice and rats has been highlighted recently (Kowash et al., 2019; Murray et al., 2019; Ozaki et al., 2020), raising concerns with which our data are in agreement. For example, the studies of maternal immune activation in rats by Kowash et al (2019) and Murray et al (2019) both report maternal plasma IL-6 increases varying from 0% to 15000%, 3h after poly I:C administration (Kowash et al., 2019; Murray et al., 2019). In our study, one dam also showed minimal plasma responses to poly I:C for some cytokines and chemokines, despite showing clear placental induction of IL-6 and IL-10. The reasons for this great variability in response, even when using a single batch/source of poly I:C, are not clear. A possibility is that the level of basal TLR3 expression in immune cells is especially sensitive to the history of prior exposure to immune stimuli (Heinz et al., 2003; Sirén et al., 2005; Ding et al., 2017), something which is not necessarily controlled in MIA studies. However, this level of variability is undesirable in an experimental context.

The elevated plasma levels of LIF after exposure to either poly I:C or resiquimod are notable, since maternal LIF is linked to the trans-placental control of foetal neurogenesis (Wright et al., 2006; Simamura et al., 2010), and to the modulation of foetal responses to elevated maternal glucocorticoids (Ware et al., 2003). Induction of maternal LIF may be important for the effects of MIA on foetal brain development and psychiatric disease risk.

### Effects in placenta

In previous MIA studies, maternal LPS administration elevated amniotic fluid Tnfα and IL-6 protein after 4h (Oskvig et al., 2012). Poly I:C administration increased Tnfα, Il-6 and Il-10 protein in placenta after 3h (Mueller et al., 2019), increased placental *Il-6* mRNA after 3h (Wu et al., 2017), and increased other cytokine mRNAs (e.g. *Tnfα Il-10*) to a much lesser degree (Hsiao and Patterson, 2011). We observed placental induction of *Il-10* but not *Tnfα* by poly I:C at 4h after administration. The slight discrepancy with respect to *Tnfα* mRNA may simply reflect the different time point. We previously reported elevated placental Cxcl10 protein 6h after poly I:C administration (Openshaw et al., 2019), and here we observed elevated *Cxcl10* mRNA levels at 4h after administration.

The effects of resiquimod in placenta were greater than those observed with poly I:C for *Il-6, Il-10* and *Cxcl10*. Induction of other immune mediators was detected after resiquimod administration, where no poly I:C effects were observed (e.g. *Ccl5, Ccl11, Cxcl11, Tnf*α mRNAs). The data show that TLR7/8 stimulation can generate a powerful immune response in placental tissue in mice. It is of interest that human umbilical cord blood cells also appear more sensitive to resiquimod than to poly I:C, in terms of induction of IL-6, IL-10 and TNFα (Στινσον ετ αλ., 2019) is true in human tissue.

From a mechanistic perspective, both a pro-inflammatory (IL-6 and TNFα) and an anti-inflammatory (IL-10) response, as well as an anti-viral (CXCL10) response, have been triggered in the placenta. The more extensive response to resiquimod includes a series of chemokines - CXCL1 and CCL2 are particularly important for recruiting inflammatory monocytes, and CCL11 is an eosinophil attractant (Sokol and Luster, 2015). hence resiquimod has induced the anticipated anti-viral reaction in the placenta. However, the extent to which these immune response mediators can access the foetal compartment is not clear in most cases (see below).

### Effects in foetal brain

In foetal brain, it has been reported that protein levels of IL-1β, IL-6, IL-10 and Tnfα were elevated at 3 or 6h after maternal poly I:C administration (Meyer et al., 2006), while another study failed to detect any increase in brain levels of IL-6 or TNFα 6h after maternal poly I:C administration (Abazyan et al., 2010). Current concerns about the reproducibility of poly I:C administration have been noted above. Maternal LPS administration, though, elevates foetal brain TNFα and IL-6 protein after 4h and 24h (Oskvig et al., 2012). In principle, the origin of these cytokines could be from the maternal side of the placenta, but a small increase (< 2 fold) in *Il-6* mRNA has also been detected in foetal brain 3h after poly I:C administration, which had normalised by 6h after administration (Wu et al., 2017). We did not detect any elevations in foetal brain cytokine mRNAs at 4h after maternal poly I:C, despite the effects seen in maternal plasma and in placenta. It is possible that small effects of poly I:C on cytokine transcription in foetal brain do occur, but are very transient, and have resolved by 4h after drug administration. However, very low relative responses to TLR3 stimulation have also been noted in human foetal tissue (Stinson et al., 2019). In contrast to the lack of effect of maternal poly I:C on foetal brain cytokines and chemokines, resiquimod produced a large (10-100 fold) stimulation of *TNF*α, *Il-6, Il-10, Ccl2, Ccl5* and *Cxcl10* expression (Figure 3). Substantial induction of *Ccl11* and *Cxcl1* mRNAs were also observed. Overall, it is clear from our data that maternal exposure to resiquimod has more substantial effects than poly I:C on the foetal CNS.

Comparing the placental and foetal brain responses, poly I:C up-regulated IL-6 and IL-10 expression in placenta, but not in brain, whereas resiquimod up-regulated these cytokines in both compartments. As noted above, there is evidence that many of the effects of MIA on the foetus (i.e. brain IL-6 and Cxcl10 induction) are mediated by maternal IL-6 acting on the placenta (Hsiao and Patterson, 2012; Wu et al., 2017), and there is clear evidence that maternal IL-6 penetrates into the foetal compartment (Zaretsky et al., 2004; Dahlgren et al., 2006). In this study, the fact that poly I:C increased maternal plasma and placental IL-6 expression, but did not affect all the mediators in the foetal brain affected by resiquimod (including IL-6 and Cxcl10) suggests that other mechanisms are also involved. Therefore, while both agents increased maternal plasma IL-6, our study suggests that not all effects on the foetal brain are mediated via maternal Il-6, and that there are qualitative differences in the foetal brain response to maternal poly I:C and resiquimod administration. One possibility to consider is that resiquimod can penetrate the placental barrier and infiltrate the foetal circulation and brain, while poly I:C cannot. There are no data available concerning the ability of resiquimod to cross the placenta in any species. However, if it can penetrate into the foetal compartment, that would usefully mirror the ability of some of the infectious agents most strongly associated with schizophrenia risk, such as *Rubella* and *Toxoplasma Gondii*, that do cross the placenta (Robbins et al., 2012) and affect the foetus directly.

There were some other interesting differences between the foetal brain response and the placental response. For example, *Ccl2* expression was elevated by resiquimod in foetal brain but not in placenta. Clearly this cannot reflect different penetration/bioavailability between the two tissue compartments, as resiquimod must pass through the placenta to access the foetal brain. We suggest that the reason is that foetal brain *Ccl2* expression is from a CNS-specific cell type rather than from immune cells, and specifically from astrocytes, which are present in the developing brain at E12.5 (Fox et al., 2004). Indeed, it has been reported that Ccl2 in the brain is produced basally by astrocytes rather than microglia (Hayashi et al., 1995; Peterson et al., 2004), and that TLR7 stimulation causes Ccl2 release from astrocytes but not from microglia (Butchi et al., 2010). Thus, the *Ccl2* mRNA induction may be demonstrating a direct or indirect action of resiquimod on foetal astrocytes.

### Evidence for microglial activation in foetal brain

The need to understand the consequences of maternal immune activation (MIA) with ss-virus mimetics becomes particularly important, considering that we have recently demonstrated that the microglial responses to stimulation of TLR3, TLR4 and TLR7 are very different (Kwon et al., 2021). While a number of reports suggest microglial activation in foetal brain following MIA in rats or mice, (Prins et al., 2018; Murray et al., 2019; Ozaki et al., 2020), there is evidence that maternal poly I:C administration does not evoke activation of foetal brain microglia (Smolders et al., 2015). We have previously found that LPS, poly I:C and resiquimod produce differing responses in microglial cells *in vitro* (Kwon et al., 2021), suggesting that distinct responses to these immune mimetics are also likely to be observed in foetal brain after MIA.

Resiquimod increased the expression of *Aif1* mRNA (encoding the widely used immune cell/ monocyte/macrophage marker Iba1)(Ribeiro et al., 2013; Elmore et al., 2014; Bruttger et al., 2015), consistent with either activation or proliferation of microglial cells. However, resiquimod decreased the expression of *Tmem119* mRNA and *Cx3cr1* mRNA, in foetal brain. *Cx3cr1*, like Iba1, is commonly used as an immune cell marker, identifying microglia, macrophages and monocytes (Harrison et al., 1998; Ginhoux et al., 2010; Mizutani et al., 2012). *Tmem119* expression has been recently characterised as being specific to microglia, and not expressed in macrophages or monocytes (Bennett et al., 2016). However, its biological functions are not fully studied yet.

Microglia respond to immune stimulation with reduced *Tmem119* expression (Bennett et al., 2016; Sousa et al., 2018; Jordão et al., 2019; Ronning et al., 2019) and reduced *Cx3cr1* expression (Boddeke et al., 1999; Wynne et al., 2010; Haimon et al., 2018; Sousa et al., 2018; Yosef et al., 2018). In fact, increased expression of *Ccl5* and *Aif1*, and decreased expression of *Tmem119* and *Cx3cr1*, appears to be a highly-characteristic signature of microglial activation (Butovsky et al., 2014; Zhang et al., 2019). Our data therefore are strongly suggestive of microglial activation in the foetal CNS, following maternal TLR7/8, but not TLR3, stimulation.

The reason for the decreased *Ccr2* expression after MIA is less clear. CCR2 is traditionally viewed as a monocyte/macrophage marker, not expressed in microglial cells (Prinz et al., 2011; Mizutani et al., 2012; Greter et al., 2015). However, the decreased expression of *Ccr2* mRNA after MIA is difficult to reconcile with altered monocyte presence in the foetal brain. There is recent evidence that microglia at E12-14 transiently express low levels of *Ccr2* (Kierdorf et al., 2013; Chen et al., 2020), which then becomes undetectable later in development. The *Ccr2* signal in the foetal brain at E12.5 may therefore derive from microglial cells. TLR4 activation by LPS reportedly decreases CCR2 expression in monocytes (Sica et al., 1997; Weber et al., 1999; Parker et al., 2004; Heesen et al., 2006; Sousa et al., 2018). We favour the interpretation that the diminished *Ccr2* mRNA expression after poly I:C and resiquimod is a similar phenomenon reflecting immune activation, but in developing microglia.

In contrast to Ccr2 and Cx3cr1, Ly6c protein levels are thought to be unchanged by immune stimuli (Jordão et al., 2019; Ronning et al., 2019). Like Ccr2, Ly6c2 is viewed as a specific monocyte/macrophage marker, not present in microglia, and this indeed appears to be the case whatever the stage of microglial development (Butovsky et al., 2014; Bowman et al., 2016; DePaula-Silva et al., 2019; Jordão et al., 2019; Ronning et al., 2019). The lack of change in *Ly6c2* expression that we observe in foetal brain after MIA suggests that there is no overt infiltration of the foetal brain by monocytes.

## Conclusions

The use of poly I:C administration to pregnant rats or mice, as a means of modelling the increased risk for schizophrenia caused by *in utero* exposure to infection, has become widespread (Zuckerman et al., 2003; Meyer et al., 2005; Estes and McAllister, 2016). Recently, however, it has become clear that variability in the characteristics of the poly I:C used are a significant issue (Kowash et al., 2019; Mueller et al., 2019). We propose that the use of resiquimod as an alternative may offer advantages in terms of reproducibility between laboratories. Furthermore, resiquimod has some conceptual advantages over poly I:C, specifically in relation to modelling schizophrenia risk. Increased risk of schizophrenia is linked with *in utero* exposure not only to influenza and rubella exposure (Mednick et al., 1988; Brown et al., 2001; Brown, 2011, 2012; Canetta and Brown, 2012), (both ss-viruses), but also to bacterial (Sorensen et al., 2009; Younga H. Lee et al., 2020) and parasitic (*Toxoplasma Gondii*) infections (Alan S. Brown et al., 2005; Brown, 2012), suggesting that the exact nature of the infectious agent is not a critical factor. However, the epidemiological evidence linking schizophrenia risk to maternal infection specifically by ds-viruses (e.g CMV, HSV-1) is weak (Khandaker et al., 2013). Conversely, the evidence for maternal ss virus infection as a risk factor is strong. Furthermore, *Toxoplasma Gondii* also activates TLR7 (Yarovinsky and Sher, 2006), as do many bacteria (Mancuso et al., 2009). It is also clear that the nature of the immune response in the brain is different to different infectious agents (Olson and Miller, 2004; Kwon et al., 2021).

Hence, until there is a more complete understanding of the mechanisms linking in utero exposure to infection with disease risk, it may be prudent to consider the use of a bacterial or ss-virus mimetic, for rodent models relating to schizophrenia. Furthermore, since resiquimod, in contrast to poly I:C, is likely to penetrate the placenta and enter the foetal compartment, it may be a much better model for agents such as influenza (Raj et al., 2014), rubella (Leung et al., 2019) and *Toxoplasma Gondii* (Robbins et al., 2012), which can also cross the placenta. A direct action of the infectious agent on the foetal brain may be essential for the disease-relevant mechanisms to be triggered. We therefore put resiquimod forward as a useful agent for modelling schizophrenia environmental risk mechanisms in rodents, to reduce experimental variability and maximise construct validity.

## List of abbreviations

ds: double-stranded
LPS: lipopolysaccharide
MFI: mean fluorescence intensity
poly I:C: polyinosinic: polycytidylic acid
ss: single-stranded
TLR: toll like receptor

## Declarations

### Ethics approval and consent to participate

In vivo experiments were performed according to Home Office (UK) regulations, and were approved by the University of Glasgow Ethical Review Board.

### Consent for publication

Not applicable

### Availability of data and materials

The datasets used and/or analysed during the current study are available from the corresponding author on reasonable request.

### Authors’ contributions

JK performed all of the experiments and contributed to an initial draft of the paper. MS and AM provided important practical contributions to many of the experiments. JC and BM supervised the study, and provided advice on data analysis and interpretation. BM conceived the study and wrote most of the manuscript.

## Acknowledgements

Not applicable

